# Perfect imperfections: seeking molecular and cellular asymmetries in the mouse brain

**DOI:** 10.1101/2025.09.11.675542

**Authors:** D.J. Houwing, Meng-Yun Wang, Andrew Silberfeld, Josephus A. van Hulten, Lukas Lütje, Judith R. Homberg, Simon E. Fisher, Joanes Grandjean, Clyde Francks

## Abstract

Although perfect symmetry is a mathematical ideal, many *Bilateria* exhibit left-right asymmetries of body, brain and behavior that are adaptive ^1-3^. In the human brain, various aspects of cognition are lateralized, including language ^4^ and handedness ^5^, which rely on left hemisphere dominance in most people. Brain hemispheric specialization can support parallel processing, neural efficiency, and action selection ^3,6^, while atypical structural or functional asymmetries are often found in neuropsychiatric disorders ^7-10^. Mice may be useful models for studying brain asymmetry because they show hemispheric differences of structure ^11,12^ and neurophysiology ^13^, but the extent to which molecular and cellular differences are involved remains unknown. We applied spatial transcriptomics with single molecule resolution to coronal sections from 31 adult mouse brains, to assess left-right differences of gene expression and cell types. Sections were chosen to capture the hippocampus and auditory cortex, as these two regions have shown the most prior evidence for functional asymmetry ^13-15^. In the hippocampus, *Crhr1* (Corticotropin-Releasing Hormone Receptor 1) was more highly expressed in the pyramidal layers of CA1, CA3 and the dentate gyrus of the left hemisphere compared to the right. In the auditory cortex, three principal components capturing transcriptomic variation showed hemispheric differences, including prominent contributions from genes that have shown associations with human brain asymmetry. Overall cellular density and cell type proportions generally showed no significant hemispheric differences, with Car3 cortical excitatory neurons showing the most laterality. The transcriptional left-right differences that we found may relate to functional asymmetries for learning, memory and hearing.

**Graphical Abstract:** 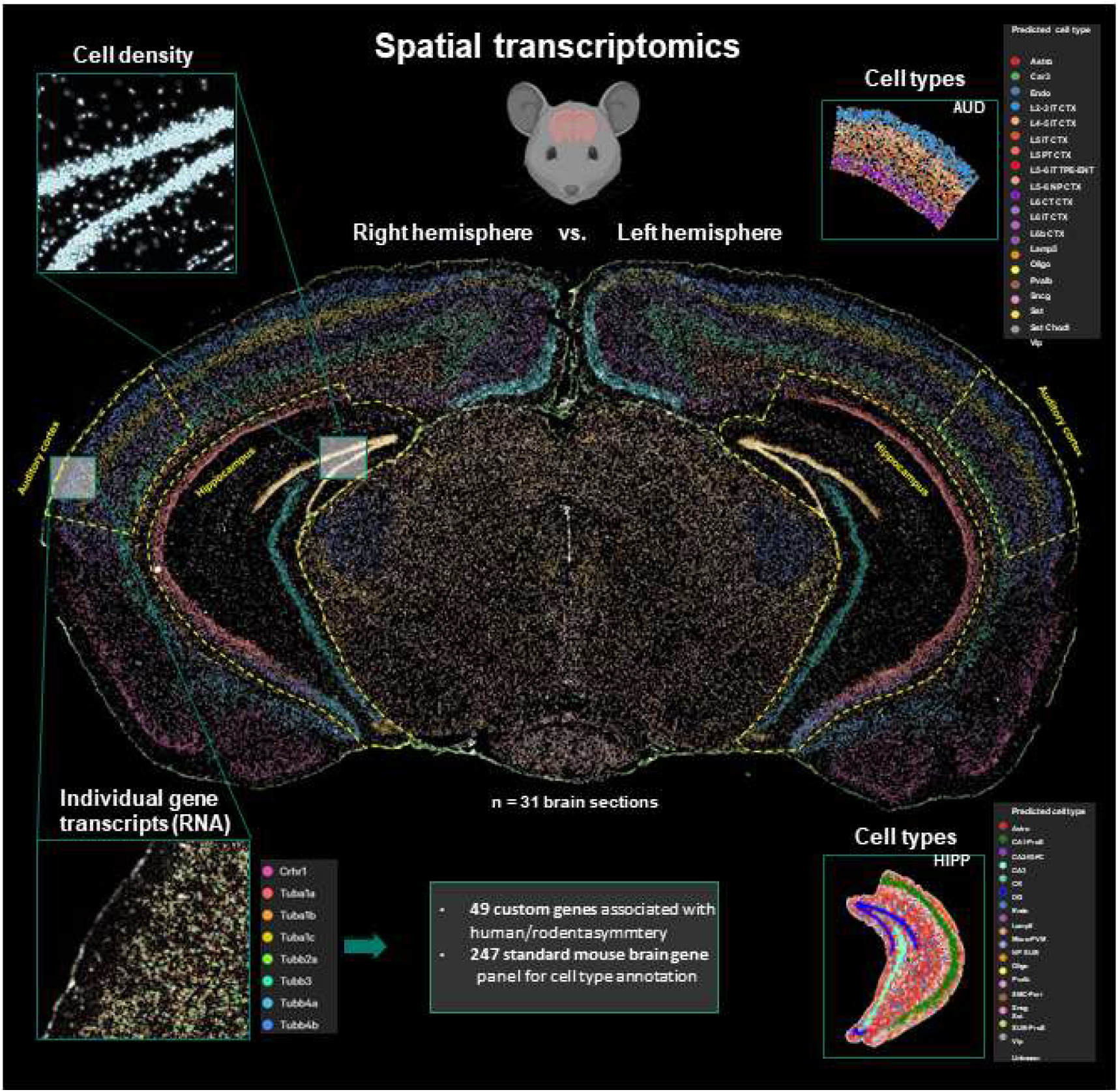

**Highlights:**

- Molecular and cellular basis of functional brain asymmetry is barely known
- Spatial transcriptomics with cellular resolution applied to mouse brain coronal sections
- Genes implicated in human brain asymmetry showed hemispheric differences in mice
- No hemispheric differences of cell type abundances

## Results and discussion

### Spatial transcriptomics to detect hemispheric differences

Previous spatial transcriptomic datasets from the adult mouse brain have not been well suited to assess left-right differences. Some have not captured both hemispheres ^16-18^, while others were based on just one or a handful of brains ^19,20^, which meant limited statistical power to detect potentially subtle asymmetries of gene expression. In addition, datasets have not generally been collected with asymmetry in mind during sectioning. For example, for coronal sections this has meant that left-right differences might be confounded with the anterior-posterior axis, whenever the cutting plane was not perpendicular to that axis.

From each of 31 adult C57BL/6J mice aged 10-16 weeks (18 males, 13 females), we subjected one coronal brain section to the Xenium spatial transcriptomics workflow (Xenium analyzer, 10x Genomics) ^21^, a method that detects single RNA molecules and places them in cellular context through their locations in three dimensions (See figure 1). After registration of the images to a reference atlas and making non-linear adjustments (QuickNII and VisuAlign: See methods) (Figure 1), we observed that sections were close to our targeted location of -3.0 Bregma (range -2.7 to -3.5) on the anterior-posterior axis (Figure S1). Our mean angle of cutting to the anterior-posterior axis was 2.03 degrees on the sagittal plane (range -14.11 to +10.63 degrees) and only -0.03 degrees on the horizontal plane (range -1.58 to +1.03) (Figure S1). The latter effectively confirmed left-right anatomical matching on average within our sections.

**Figure 1.**
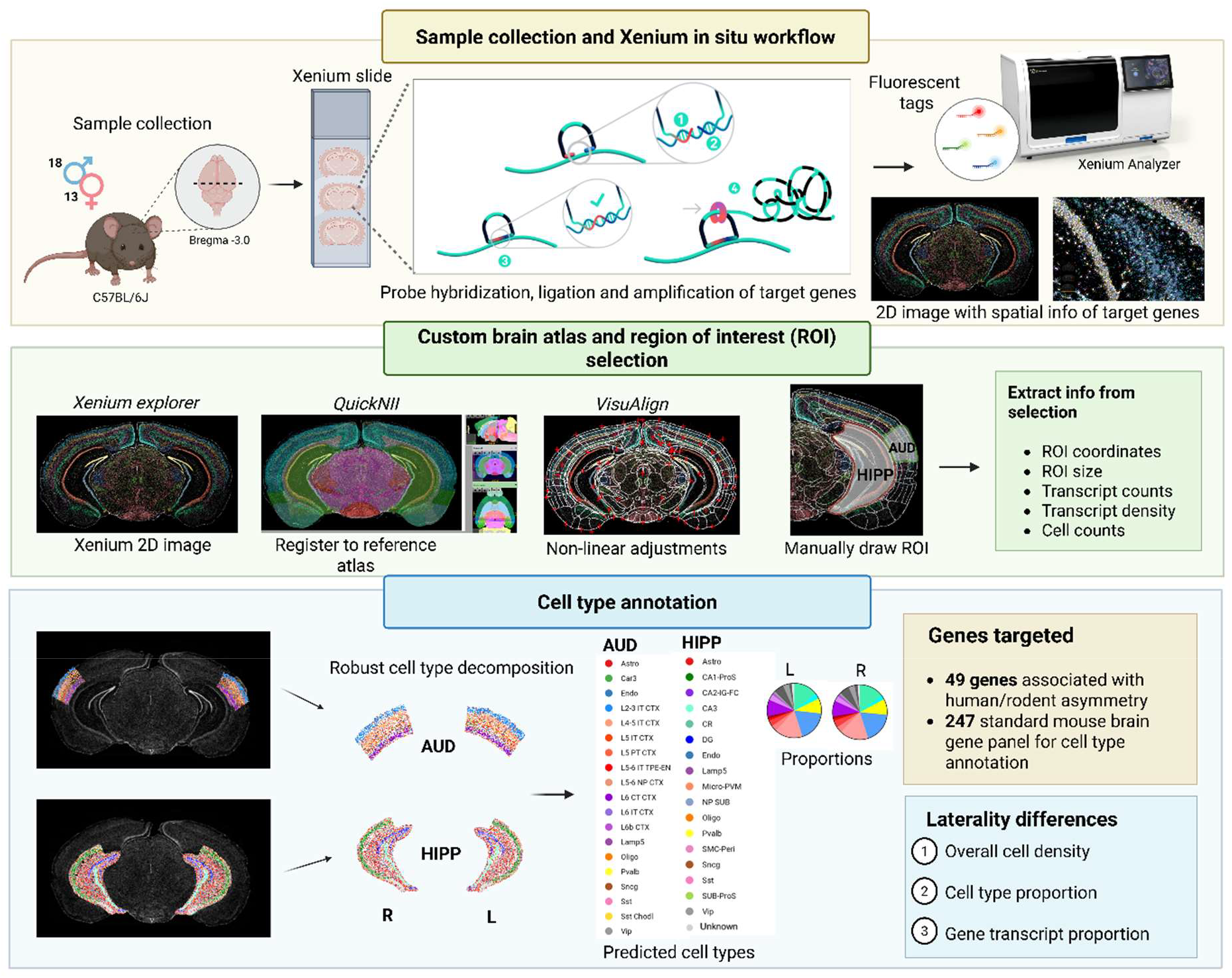
Schematic of the study workflow. Coronal brain sections were extracted from wild-type adult mice and subjected to Xenium spatial transcriptomics, a technique for detecting single transcript molecules with subcellular spatial precision. Images were aligned and adjusted to an atlas for region-of-interest identification and data extraction (transcript and cellular statistics). Major cell types were annotated based on the application of single cell RNA sequencing data. Hemispheric differences of transcript and cellular abundance were investigated. AUD: auditory cortex. HIPP: hippocampus. L: left. R: right.

The gene panel consisted of a standard Xenium panel of 247 genes selected to distinguish major cell types and/or with key functions in the mouse brain, plus a custom set of 49 genes that included 31 orthologs of genes implicated by large-scale genome-wide association studies of human brain asymmetry ^22^ and handedness ^23-26^, as well as genes implicated in brain or body asymmetry in mice or flies ^27-30^, and genes involved in neurotransmission that might support hemispheric functional specialization at the synapse (see Table S1 for summary information on all 49 custom genes). We detected a mean of almost 120.000 cells and nearly 70 million reads per section, with a median of 321 transcripts per cell.

We extracted data from two regions of interest per hemisphere - the hippocampus and auditory cortex (Methods; Figure 1). These two regions have shown the most extensive evidence for functional asymmetry of any rodent brain structures, based on neurophysiological and behavioural data ^13-15^ (discussed in more detail in the two sections directly below). Four sections were omitted from analysis of the hippocampus, and eleven from the auditory cortex analysis, due to damage or folding in these regions. This yielded sample sizes of 28 sections for hippocampus and 21 sections for auditory cortex with usable paired left-right data. Per region and hemisphere, we extracted the region coordinates, transcript counts per gene, transcript densities per gene, and total cell counts.

Through use of mouse hippocampus and whole cortex single cell RNAseq datasets ^31^, we then annotated cells in our Xenium data to 17 major cell types for the hippocampus and 19 for the auditory cortex from each section and hemisphere, based on clustering through means of differential gene expression profiles (Methods). Annotated cell types included major excitatory and inhibitory neuronal classes and glial cells, such as CA1, CA3 and dentate gyrus (DG) pyramidal neurons, and cortical layer-specific cell types (Figures 1, 2, 3).

**Figure 2.**
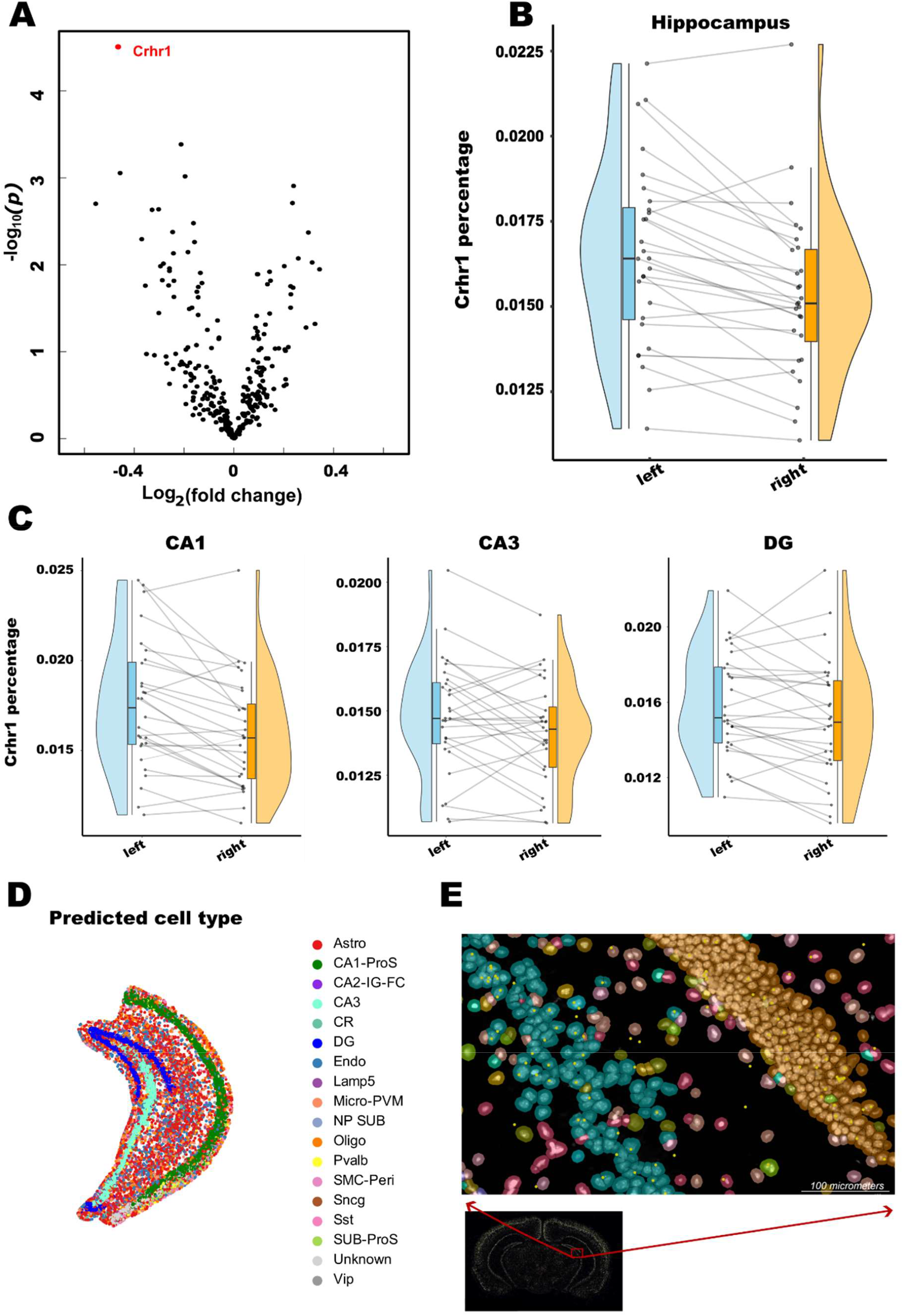
Examining molecular and cellular asymmetries in the mouse hippocampus. **A** Volcano plot of gene expression in the hippocampus (N=28 animals). Each dot denotes a gene. The x-axis represents the log_2_ fold change in the right hemisphere compared to the left hemisphere. The y-axis depicts the p values. The red dot indicates Crhr1 (t=-4.90; P=3.1×10^-5^, FDR-adjusted P=0.0092). **B** Crhr1 expression in the left and right hippocampus. The x-axis indicates the hemisphere while the y-axis represents Crhr1 expression as a percentage of all transcripts in the ROI of a given hemisphere. Left and right sample pairs (dots) from the same animal are linked by lines. Box plots show the median, 25^th^ and 75^th^ centiles, with whiskers indicating 1.5 times these centiles. Density plots are also shown. **C** Crhr1 expression in left and right CA1, CA3 and DG subfields respectively. **D** Cell type annotation of the hippocampus in an example section and hemisphere (sample F687, left hemisphere). Note that only 15 cell types were abundant enough for robust testing of hemispheric differences (see Methods and Table S2). Astro: Astrocytes; CA1-ProS: Neurons in the CA1 region of the hippocampus and prosubiculum; CA2-IG-FC: neurons in the CA2 region of the hippocampus; CA3: neurons in the CA3 region of the hippocampus; CR: Cajal-Retzius cell type; DG: neurons in the DG region of the hippocampus; Endo: Endothelial cells; Lamp5: Lysosome-associated membrane protein 5-expressing inhibitory neurons; Micro-PVM: Microglia and perivascular macrophages; NP SUB: Near-projecting, subiculum. Oligo: Oligodendrocytes; Pvalb: Parvalbumin-expressing inhibitory interneurons; SMC-Peri: Smooth Muscle Cell / Pericyte; Sncg: Synuclein gamma-expressing inhibitory neurons; Sst: Somatostatin-expressing inhibitory neurons; SUB-ProS: Neurons in the subiculum and prosubiculum; Vip: Vasoactive intestinal peptide-expressing inhibitory neurons. **E** Crhr1 expression in an example section, and locations of Crhr1 transcripts in relation to cell bodies of DG and CA3. Note that this panel is extracted from the Xenium software that uses unsupervised clustering for cell types rather than annotating pre-defined types based on single cell RNA sequencing data. The most abundant types visible are DG excitatory (gold) and CA3 pyramidal (blue) cells. Green dots show individual Crhr1 transcripts.

**Figure 3.**
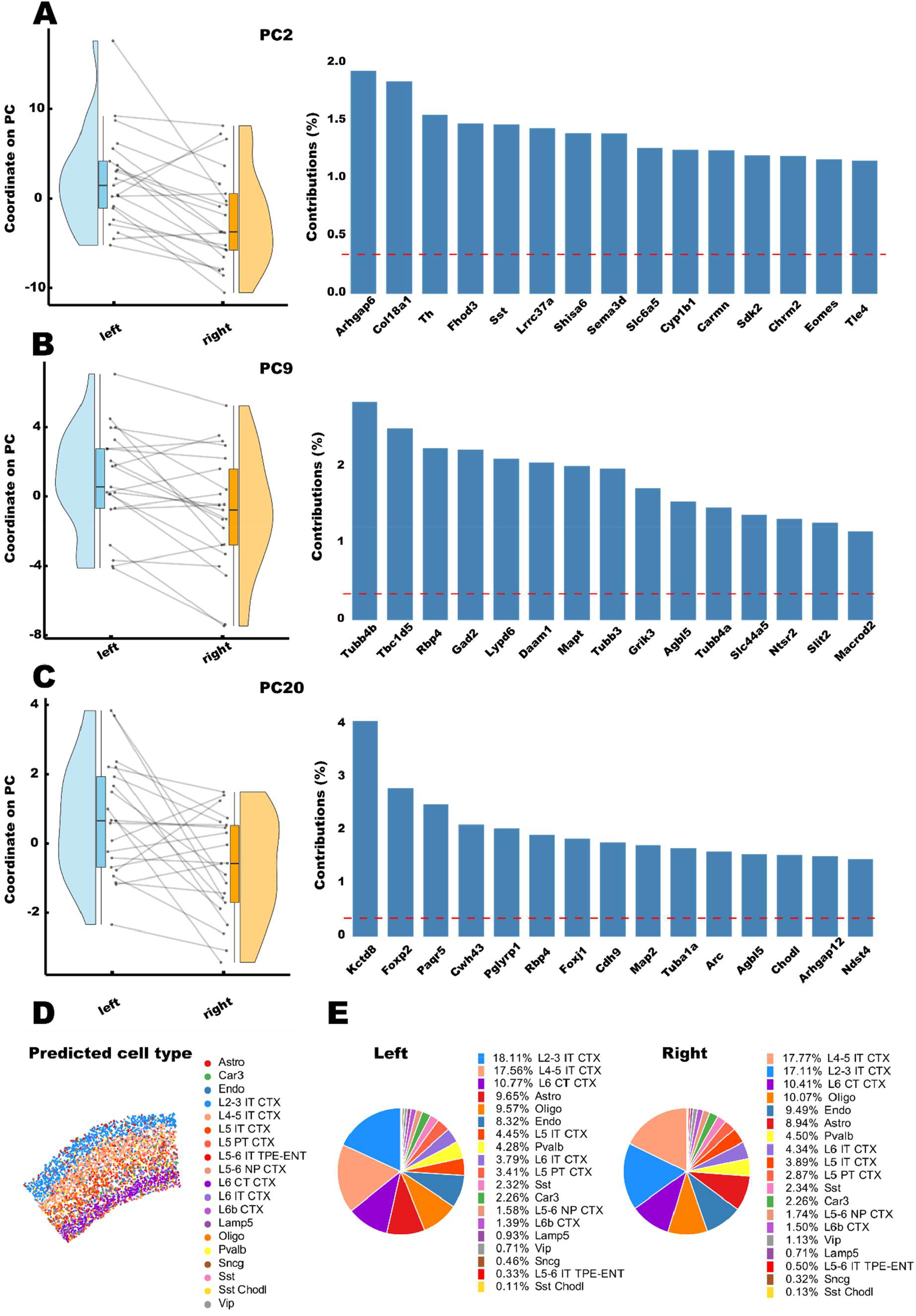
Examining molecular and cellular asymmetries in the mouse auditory cortex. N=21 animals. Three principal components of transcriptional variance in the auditory cortex were linked to hemispheric differences: ***A*** the second, ***B*** the ninth and ***C*** the twentieth components. In the left-side panels the x-axis indicates the hemisphere while the y-axis represents sample coordinates along a given PC. Left and right sample pairs (dots) from the same animal are linked by lines. Box plots show the median, 25^th^ and 75^th^ centiles, with whiskers indicating 1.5 times these centiles. Density plots are also shown. In right-side panels, the loadings of the top 15 individual genes on the principal components are shown (dashed horizontal lines show the average loadings across 296 genes). ***D*** Cell type annotation in the auditory cortex of an example section and hemisphere (sample F087, left hemisphere). Note that only 15 cell types were abundant enough for robust testing of hemispheric differences (see Methods and Table S2). ***E*** Pie charts of cell type percentages in the left and right auditory cortex. Astro: Astrocytes; Car3: Carbonic anhydrase 3-expressing neurons; Endo: Endothelial cells; L2-3 IT CTX: Layer 2/3 intratelencephalic cortical neurons; L4-5 IT CTX: Layer 4/5 intratelencephalic cortical neurons; L5 IT CTX: Layer 5 intratelencephalic cortical neurons; L5 PT CTX: Layer 5 pyramidal tract cortical neurons (involved in motor control); L5-6 IT TPE-ENT: Layer 5/6 intratelencephalic neurons in the temporal and entorhinal cortex; L5-6 NP CTX: Layer 5/6 near-projecting neurons in the cortex; L6 CT CTX: Layer 6 corticothalamic neurons; L6 IT CTX: Layer 6 intratelencephalic cortical neurons; L6b CTX: Layer 6b cortical neurons; Lamp5: Lysosome-associated membrane protein 5-expressing inhibitory neurons; Oligo: Oligodendrocytes; Pvalb: Parvalbumin-expressing inhibitory interneurons; Sncg: Synuclein gamma-expressing inhibitory neurons; Sst: Somatostatin-expressing inhibitory neurons; Sst Chodl: Somatostatin- and Chodl-expressing inhibitory neurons. Vip: Vasoactive intestinal peptide-expressing inhibitory neurons.

With these data we were able to test for hemispheric differences of transcript abundances, in terms of individual genes and also principal components capturing major dimensions of transcriptome variation, as well as total cell densities, and major cell type proportions (Figure 1) (see Methods for the statistical approaches). Rarer cell types in each region were excluded (Methods) from testing for hemispheric differences, which left 15 cell types in the hippocampus and 15 in the auditory cortex (Table S2)). We make our full Xenium dataset freely available for download (see the ‘Resource availability’ section).

### Asymmetries in the hippocampus

The mouse hippocampus has been reported to be lateralized with respect to spatial learning and memory ^14,15^. The left hemisphere preferentially stores spatial information as discrete, salient locations, while the right hemisphere represents space more continuously. This laterality is thought to contribute to route computation and flexible navigation ^14,15^. Other work has indicated that in the hippocampal DG, higher context discrimination occurs in the left hemisphere, while there is a bias towards generalization in the right hemisphere ^32^.

By screening all 296 genes in our Xenium data for hemispheric differences in the hippocampus (Figure 2, Table S3) we identified one gene as asymmetrically expressed at false discovery rate (FDR) < 0.05: Crhr1 was more highly expressed in the left hippocampus compared to the right hippocampus (t = - 4.90; P = 3.1×10^-5^, FDR-adjusted P=0.0092) (Figure 2). Post hoc analysis showed this asymmetry to be present in each of the subfields CA1 (t=-4.33, P=0.00015), CA3 (t=-2.26, P=0.031), and DG (t=-2.31, P=0.028) (Table S4), with the strongest effect in CA1 where the level of Crhr1 was 1.38 times higher on the left than the right (Figure 2). Crhr1 transcripts predominantly co-localized with the pyramidal layers of CA1 and CA3 and the granule cells of DG, which are comprised mostly of excitatory neuronal cell bodies (Figure 2).

Corticotropin-releasing hormone signaling contributes to diverse stress-related functions in the mammalian brain, including motivation and emotion ^33^. Prenatally stressed mice show hippocampal methylation and expression changes of Crhr1 ^34^, while neonatal stress affects hippocampal neuron morphology and CRHR1 protein levels ^35^. In addition, spatial learning and memory deficits induced by prenatal glucocorticoid exposure depend on hippocampal CRHR1 signaling ^36^. Crhr1 gene expression level, in the left hippocampus specifically, has also been linked to prenatal stress in the contactin-associated protein-like 2 (Cntnap2) mouse model for autism spectrum disorder ^37^. We included Crhr1 in the current study because large-scale genome-wide association analysis in humans has implicated the *CRHR1* locus in human structural brain asymmetry ^22^, including asymmetry of hippocampus volume and parahippocampal cortex surface area, as well as left-handedness ^25^. Taking these various observations together with the current study, Crhr1 is a strong candidate for affecting hemispheric specialization of neuronal functions in the hippocampus.

The estimated proportion of true nulls (Methods) among the 296 genes was 63% when testing for hemispheric differences in the hippocampus, so that some of the nominally significant genes that did not surpass multiple testing correction, with asymmetries less strong than Crhr1 (Table S3), may also be asymmetrically expressed to subtle extents. We applied principal component analysis that identified 18 components capturing 90% of transcriptome variance in the hippocampus, across sections and hemispheres (Methods) (Figures S2, S3). None of these components showed significant hemispheric differences (all FDR adjusted P>0.32), i.e. data sampled from the left versus right hippocampus did not differ significantly in their coordinates along any of the 18 principal components (Table S5). Functional laterality of the hippocampus may therefore involve left-right differences in the expression levels of a minority of genes from different co-expression networks, rather than being reflected in major components of transcriptional variance.

An asymmetry in hippocampal cellular morphology has been reported at the synaptic level: hippocampal CA1 pyramidal cell synapses differ in size and shape depending on whether they receive input from the left or right CA3 ^28^. At the molecular level, asymmetric synaptic distribution of glutamate receptor subunits was reported in Schaffer-collateral CA1 pyramidal cells ^27-29^. Asymmetries in hippocampal synapses are accompanied by asymmetry in neuronal plasticity, measured by long term potentiation differences ^27-29^. For glutamate receptor subunits specifically, our Xenium data included Grin2a, Grin2b and Gria1 (encoding NR2A, NR2B and GLUR1 respectively). We saw no significant asymmetries of expression for these three genes in the hippocampus, nor in subfields CA1, CA3 or DG when tested separately in post hoc analysis (all P>0.05) (Tables S3, S4), although receptor subunit proportions at the whole structure or subfield level are not necessarily indicative of specific synaptic subtype abundances or sizes ^28^ .

Overall cell density in the hippocampus showed no significant hemispheric difference (t=1.08, P=0.29, Table S6), and neither did the proportion of any of 15 major hippocampal cell types (all FDR adjusted P>0.35), including subfield-specific neurons of CA1, CA3 and DG (Table S7, Figure S4). Therefore, the mouse hippocampus appears remarkably symmetrical in terms of total cellular and cell type abundances, considering the functional laterality ascribed to this brain structure. Functional laterality in the hippocampus may therefore be supported by asymmetrical expression of a minority of genes expressed from cell types that do not differ in their abundances between hemispheres.

### Asymmetries in the auditory cortex

The mouse auditory cortex plays a crucial role in perceiving and processing auditory information, and does so in a lateralized manner. The left auditory cortex of adult female mice is dominant for the processing of pup ultrasonic vocalizations, which is crucial for maternal behavior like pup recognition and retrieval ^38-40^. The right auditory cortex, in contrast, is more engaged for neural and behavioral responses to frequency sweeps (sounds that change in frequency over time) ^39^, as well as echoic memory (a type of sensory memory that briefly stores auditory information, acting as a temporary buffer for sounds) ^41^. During brain development, maturation of auditory cortex circuits in the left and right hemispheres occurs asynchronously, with the right maturing earlier than the left ^13^.

When testing all 296 genes in our Xenium data for hemispheric differences in the auditory cortex (Figure S5, Table S8), we identified no individual genes with significant asymmetrical expression after FDR adjustment (all FDR adjusted P>0.061). However, Crhr1 was among the genes that showed nominally significant asymmetry (i.e. before FDR adjustment) in the auditory cortex (t=-3.4, P=0.0025; Table S8), and the estimated proportion of true nulls among the 296 genes (Methods) was 47%, suggesting that some of the nominally significant asymmetries were real in this structure.

Principal component analysis identified 20 components that together captured 90% of transcriptome variance in the auditory cortex, across sections and hemispheres (Methods; Figures S6, S7)). Unlike the hippocampus where no principal components reflected hemispheric differences, left and right data from the auditory cortex differed significantly in their coordinates along three principal components with FDR-adjusted P<0.05 (Figure 3-; Table S9): component 2 (t=-3.68, P=0.0015) captured 11.1% of overall transcriptome variance with strongest loadings from the genes Arhgap6 and Col18a1 (Figure 3; Table S10); component 9 (t=3.38, P=0.003) captured 3.6% of overall variance with microtubule-related genes Tubb4a, Tubb4b, Tubb3, Daam1, Mapt and Agbl5 among the main contributors (Figure 3; Table S10); component 20 (t=2.99, P=0.007) captured 1.0% of overall variance with the strongest loadings from genes Kctd8 and Foxp2 (Figure 3; table S10). Interestingly, *COL18A1* (collagen type XVIII alpha 1 chain) and microtubule-related genes *TUBB4A, TUBB4B, TUBB3, DAAM1, MAPT* and *AGBL5* were all found in large-scale genetic studies in humans to be associated with structural brain asymmetries ^22,24,26^ and/or handedness ^25^. In addition, loss of function variants in human *FOXP2* cause a severe speech and language disorder ^42^, while common single nucleotide polymorphisms in this gene were associated with functional hemispheric asymmetry for speech perception in one candidate gene study ^43^. Principal components 2, 9 and 20 together comprised 15.7% of overall transcriptome variance, so that their hemispheric differences suggest a more pervasive asymmetry of gene expression in the auditory cortex than in the hippocampus.

There was no significant asymmetry of total cell density in the auditory cortex (t=1.31, P=0.20, Table S6). Likewise, when considering the proportions of 15 major cortical cell types, including layer-specific excitatory and inhibitory cell types (Figure 3), no significant asymmetry was detected in the auditory cortex (all FDR adjusted P>0.12) (Table S11). Car3 neurons showed the most nominally significant asymmetry (i.e. prior to FDR correction: t=-2.75, P=0.008), indicating a higher proportion in the left auditory cortex than the right. These excitatory neurons of the deep cortical layers have a relatively limited intracortical projection pattern ^44^, so that a higher proportion in the left auditory cortex may enhance local rather than global information processing on that side.

A previous study of the mouse brain based on serial two-photon tomography images (i.e. without molecular resolution of cell types) reported extensive hemispheric differences of total cellular density for many brain regions ^45^. In that study, for the auditory cortex, a cellular density difference of more than 20% was reported between left and right hemispheres in layers 2/3. Although the sample size of that study was large (N=507), it was based on previously published data from a connectome study that involved unilateral surgery, injection of recombinant adeno-associated virus and iontophoresis into the right hemisphere specifically, followed by some weeks of recovery prior to euthanasia and imaging ^46^. It seems possible that laterality may have been affected by this procedure. In addition, study-specific technical factors during automated image registration may affect estimates of regional volume asymmetries^12^, which in turn would affect cellular density measurements. The present study indicates that, like the hippocampus, the auditory cortex is remarkably symmetrical in terms of total cell density and most major layer-specific cell types, given the developmental and functional laterality reported for this region. Our data suggest instead that hemispheric differences of transcriptome profiles are more important for functional laterality of the mouse auditory cortex than hemispheric differences of total cell density or cell type proportions.

### Summary and outlook

This study identified transcriptomic differences between left and right hemispheres of the adult mouse brain that may relate to functional asymmetries of learning, memory and auditory processing. Corticotropin receptor signaling was implicated in the hippocampus, while broader transcriptomic profiles were implicated in the auditory cortex, including a principal component featuring microtubule-related genes that affect human brain asymmetry and handedness. Overall cell density and cell type abundances were remarkably similar between hemispheres, although there was a suggestive asymmetry of Car3 deep-layer cortical excitatory neurons in the auditory cortex.

With appropriate multiple testing correction this study was able to detect roughly 1.4-fold differences of individual transcript levels between hemispheres. However, some functional asymmetries might be supported by even more subtle hemispheric differences of molecular abundance that affect fine tuning of neurophysiology. Therefore larger-scale studies, as well as studies of protein-level asymmetries ^28^, may be beneficial to reveal further functionally relevant asymmetries.

We focused on the hippocampus and auditory cortex on the basis of previous literature on lateralized functions, and because both regions could be captured in single coronal sections. We make our dataset available to the community (see Resource Availability) to explore additional regions in these sections. Future studies of asymmetry would benefit from serial coronal sectioning per brain along the anterior-posterior axis to capture more regions, and help address any subtle ‘cerebral torque’ phenotype whereby one hemisphere may be shifted relative to the other on the anterior-posterior axis^11,12^.

When testing for hemispheric differences we modelled a random effect of section to account for technical variation (batch, slide, Bregma position, cutting angles in different planes – see Methods) and also biological variation (e.g. age, sex, brain size, as there was one section per animal). In humans, various regional structural asymmetries in the brain can differ slightly on average between males and females ^47^. However, the present study was not designed to investigate sex effects, as sex was confounded by technical variables such as batch and slide. To investigate possible sex*hemisphere interactions while correcting for technical and biological variables as nested fixed effects, a larger sample size would be needed for robust model fitting and adequate statistical power with FDR correction. Nonetheless, our inclusion of both males and females likely adds to generalizability of our findings on hemispheric differences. For example, in *post hoc* analysis of Crhr1 in the hippocampus, nominally significant asymmetry was found separately in both females (t=-2.51, P=0.025) and males (t=-4.49, P=0.0003).

Our study was designed to detect molecular and cellular asymmetries that may contribute to hemispheric differences of neurophysiology and function in the adult mouse brain. These genes may only partly overlap with those required to pattern asymmetry during neurodevelopment. Future studies would benefit from capturing developmental trajectories of molecular and cellular asymmetries from embryo to adulthood.

## Resource availability

### Lead contact

Further information and inquiries should be directed to the lead contact, Clyde Francks.

### Materials availability

This study did not generate new unique reagents.

### Data and code availability

The dataset will be made freely available to download upon publication of this article. Code: https://github.com/MengYunWang/Mouse_brain_asym

## Acknowledgments

This research was funded by the Max Planck Society (Germany). We would like to thank the coordinators of the translational neuroscience unit (TNU) of the Donders Institute, Radboud University Nijmegen, and the animal technicians and caretakers of the animal research facility (CDL) of Radboud University Medical Center for technical support and creative input regarding animals and procedures. Thanks also to Sabrina van Heukelum for helping to set up this research line.

## Author contributions

Conceptualization: DJH, MYW, LL, SEF, JG, CF.

Methodology: DJH, MYW.

Software: DJH, MYW.

Formal analysis: DJH, MYW.

Investigation: DJH, MYW, CF.

Resources: CF.

Data curation: DJH, MYW.

Writing – original draft: DJH, MYW, CF.

Writing - Review & Editing: DJH, MYW, AS, JvH, LL, JRH, SEF, JG, CF.

Visualization: DJH, MYW.

Project administration: DJH, CF.

Funding acquisition: SEF, CF.

## Declaration of interests

The authors report no conflicting interests.

## STAR★Methods

### Key resources table

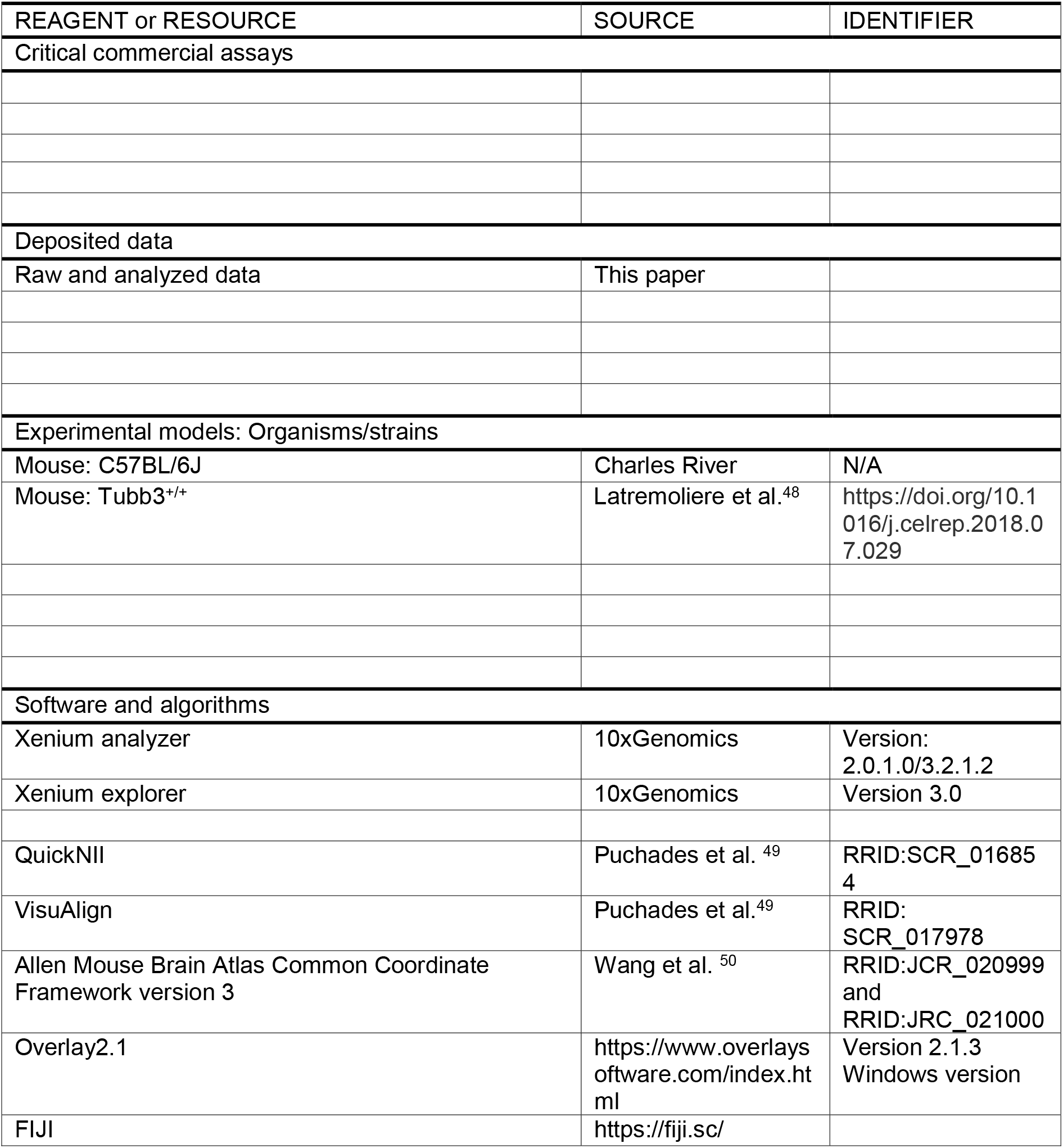
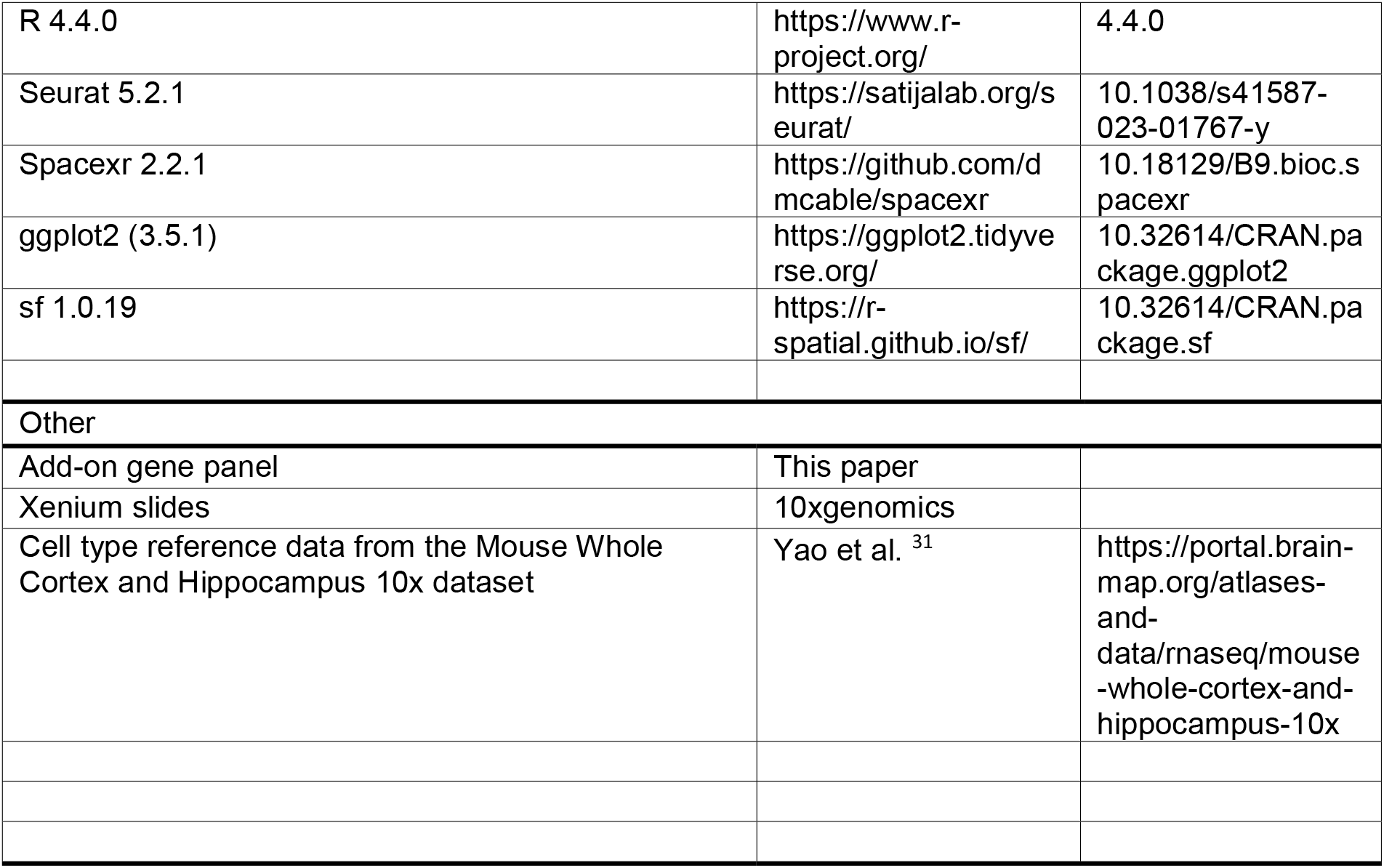

## EXPERIMENTAL MODEL AND SUBJECT DETAILS

### Mice

All experimental procedures were approved by the Central Committee for Animal Experiments, Den Haag, The Netherlands. Experiments occurred in two batches: the first batch of animals originating from Charles River France (10 males, 10 females, C57BL/6J), the second batch originating from the Radboud University breeding farm (8 males, 4 females, Tubb3^+/+^, i.e. wild-type mice from a Tubb3 knockout line ^48^). Adult male and female mice arrived at 10-16 weeks of age and were housed at the Central Animal Facility of the Radboud University Nijmegen. Mice were housed in same-sex groups (3-4 per cage) under a 12h:12h light/dark cycle (21:00 lights off) and were provided with food and water *ad libitum*. After one week of acclimatization, animals were euthanized by cervical dislocation followed by rapid brain removal. Brains were rinsed in saline and the cerebellum was removed for optimal placement in the cryomold. The brain was placed on a drop of optimal cutting temperature (OCT) compound in a pre-chilled cryomold (22 x 22 x 20mm) with the olfactory bulb facing upwards. Next, the cryomold was completely filled with OCT compound and placed on dry ice for freezing. Fresh frozen brains were stored in sealed plastic bags in -70°C.

## METHOD DETAILS

### Sample collection

Brains were sliced into coronal sections using a cryostat (Leica CM1950, Leica Biosystems, Deer Park, IL, USA) at 10 µm thickness. For each animal, one section was collected between anterior-posterior coordinates -2.92mm and -3.52mm relative to Bregma which captures both the hippocampus and auditory cortex. During cutting, left-right anatomical symmetry of the section was visually checked while advancing towards the desired Bregma coordinates. A section was collected onto a superfrost microscope slide and fixed (4% PFA, 60 sec.), washed (PBS 1x, 60sec), stained (0.1% Cresyl Violet, 60sec) and rinsed with ethanol (70% EtOH, 60sec) before again assessing symmetry visually under a brightfield microscope (Leica Microsystems). We used landmarks to determine when to collect the section according to Bregma target -3.0: after elongation of the hippocampus, the CA1 and CA3 pyramidal layers separate all the way down to the ventral hippocampus, and the corpus callosum should be disconnected between hemispheres. When both conditions were met in both hemispheres, the 5^th^ section was collected onto a Xenium microscope slide. On each Xenium slide, up to 3 sections (from 3 different animals) were placed within the imageable capture area (12mm x 24mm), while taking caution not to cover the fiducial markers on the slide. Each Xenium slide was kept in its own slide mailer and stored at -70°C. All brains were sectioned, and all sections were collected, by the same person.

#### Xenium sample preparation and on-instrument analysis

Samples were personally delivered, while kept frozen on dry ice, to the Leiden Genome Technology Center (Leiden, the Netherlands). Here, samples were prepared (Demonstrated Protocol CG000581, 10x Genomics) and subjected to the Xenium assay workflow (User Guide CG000582, 10xGenomics), followed by on-instrument pre-processing with the Xenium analyzer software (User Guide CG000584, 10xGenomics).

The Xenium data consisted of spatial information on 31 mouse brain coronal sections representing an average of almost 70 million reads and roughly 120,000 cells per brain section. (The section from a 32nd brain – female - was omitted immediately after observing an extreme cutting angle in the sagittal plane). For every transcript read, the gene identity, 3-dimensional position (x,y,z), and phred-based quality value (qv) was obtained. On average, 80% (range 76% - 84%) of the reads were of high quality (qv>20). In addition, cell-segmentation masks and a cell-by-gene matrix containing the expression and position (x,y) of every detected cell were obtained. A median of 321 transcripts were detected per cell.

### Data processing

#### Selection criteria of sections

Xenium analyzer software outputs for 31 brain sections were visualized and processed using the Xenium explorer software (10x Genomics Xenium explorer v2). Sections were visually inspected for several inclusion criteria: If the auditory cortex of one hemisphere was outside of the capture area, the section was omitted for auditory cortex analysis (but not for hippocampal analysis); if sections were damaged or folded within the auditory cortex or hippocampus, sections were omitted for their respective area analysis; if sections appeared to be visibly asymmetrical (left vs. right) anatomically (judged by looking at the hippocampus), they were omitted as well. In the end, 6 sections of the male brains were omitted for the auditory cortex (2 partially outside of the capture area, 3 due to damage, 1 due to folding) and 4 sections of the female brains (2 due to damage, 2 due to folding). For the hippocampus, 2 male sections (due to damage) and 1 female section (due to damage) were omitted.

For subsequent analysis we therefore had the following numbers with usable bilateral data: auditory cortex: 12 male, 9 female; hippocampus: 16 male, 12 female.

#### Custom atlas maps, ROI selection and data extraction

Accurately assigning the anatomical locations of our regions of interest to the images generated by Xenium analyzer was of crucial importance. Using the registration tool QuickNII ^49^, the Xenium image was registered to the Allen Mouse Brain Atlas to produce a customized atlas map that was corrected for variation in the cutting plane and overall sizes of the sections. Further nonlinear refinements were made using VisuAlign ^49^. This process yielded measures of the cutting angles for each section, through which we could verify that our sections were symmetrical on average (-0.03 degrees on the horizontal plane (range -1.58 to +1.03) (Figure S1).

Using FIJI ^51^, an image was then created of the atlas line art for each section. Using the software Overlay2.1, the atlas line art was projected on top of the Xenium image in Xenium explorer. This way, we could manually trace the exact atlas-defined regions of interest in Xenium explorer to extract data solely from our ROIs, uniquely specified for each section. Since the outer layer of the cortex was easily damaged during sectioning and mounting, we excluded layer 1 of the auditory cortex in our analysis. Extracted data included ROI coordinates, ROI size, gene transcript counts, gene transcript density and cell counts.

## QUANTIFICATION AND STATISTICAL ANALYSIS

All data processing and analysis was based on R4.4.0 (x86-64-pc-linux-gnu), where several generic libraries were used including ggplot2 (3.5.1), future (1.34.0), dplyr(1.1.4), tidyr (1.3.1).

### Gene expression asymmetry

The gene expression data of each gene was normalized by dividing by the total gene density within each section, hemisphere and region of interest. Then, the values were standardized to unit variance and zero mean, which ensured that each gene contributed equally to downstream analysis regardless of its original dynamic range. A design matrix was \ constructed with a factor hemi encoding left versus right hemisphere and a mouse-ID factor to model between-section intercepts. The mouse-ID factor therefore accounted for technical factors such as batch and slide, and also biological factors such as sex, for the purpose of assessing left-right differences. We fitted this linear model using the limma (3.60.6) ^52^ (package’s *lmFit* function which computes ordinary least squares estimates of the hemisphere coefficient while accounting for repeated measures (left and right from the same section). Finally, we applied empirical Bayes shrinkage (*eBayes* function) to obtain moderated statistics for the left–right contrast. Multiple comparison was corrected with FDR. The estimated proportion of true nulls among the 296 genes was estimated from the vector of 296 p-values using the *propTrueNull* function in limma (‘*hist*’ option), which assumes that true null p-values are uniformly distributed between 0 and 1, while non-null p-values are concentrated at lower values.

#### Principal component analysis

Given the relatively large number of 296 genes and their correlations (Figures S2, S6), data reduction was done with principal component (PC) analysis. For this purpose the data were first harmonized with ComBat ^53^ from the library *sva* (3.52.0) ^54^ to adjust for batch and other technical effects. Then, PC analysis was run using the *prcomp* function from *factoextra* library (1.0.7). The number of PCs was decided based on the accumulated explained variance which was set to a maximum 90%, which required 20 PCs in hippocampus and 18 PCs in auditory cortex. Finally, paired t tests were used to test the laterality effect, where FDR was used to correct multiple comparisons. This analysis tested whether data sampled from the left versus right hemisphere differed in their coordinates along any of the PCs.

#### Overall cell density

The overall cell density of each region within a given section and hemisphere was given by the total cell count divided by the area of the region. To test the left-right contrast, a linear model was used, with hemisphere as the fixed factor and mouse-ID as random factor.

#### Cell type density

Per-transcript location data, the cell by gene matrix, cell segmentation, and cell centroid information were loaded and processed with default parameters in Seurat (5.2.1) ^55^, which involves quality control (e.g. cells with fewer than 100 unique molecular identifiers were removed), normalization and harmonization across samples. Cell type annotation was done with robust cell type decomposition (RCTD) via the spacexr (2.2.1) library ^56^ using default parameters. RCTD is a computational method that leverages cell type profiles from a reference dataset to decompose cell type mixtures and assign cell types in a target dataset. Cell type references for the hippocampus and auditory cortex were constructed from the Mouse Whole Cortex and Hippocampus 10x dataset ^31^. RCTD required each cell to have at least 10 reads for relevant informative genes for a given cell type.

We excluded cell types with average counts across sections and hemispheres less than 30, which left 15 annotated cell types in the hippocampus and 15 in the auditory cortex (Figures 2&3; Table S2). The cell type data were then normalized for each section, hemisphere and region by dividing by the total number of annotated cells for that section/hemisphere/region. The left-right contrast was tested using a linear model with hemisphere as the fixed factor and mouse-ID as random factor, again using limma (3.60.6) ^52^ and correcting for multiple comparisons with FDR.

## Supplemental information

Document S1. Figures S1–S7

Tables S1-S11 as separate worksheets in one supplementary Excel file.

## Notes

### Competing Interest Statement

The authors have declared no competing interest.

