## Supplementary figures for "Perfect imperfections: seeking molecular and cellular asymmetries in the mouse brain"

**Contents:**

Supplementary figures S1 – S7 on pages 2-8.

(Supplementary Tables S1-S11 are given as separate worksheets in an accompanying Excel file.)

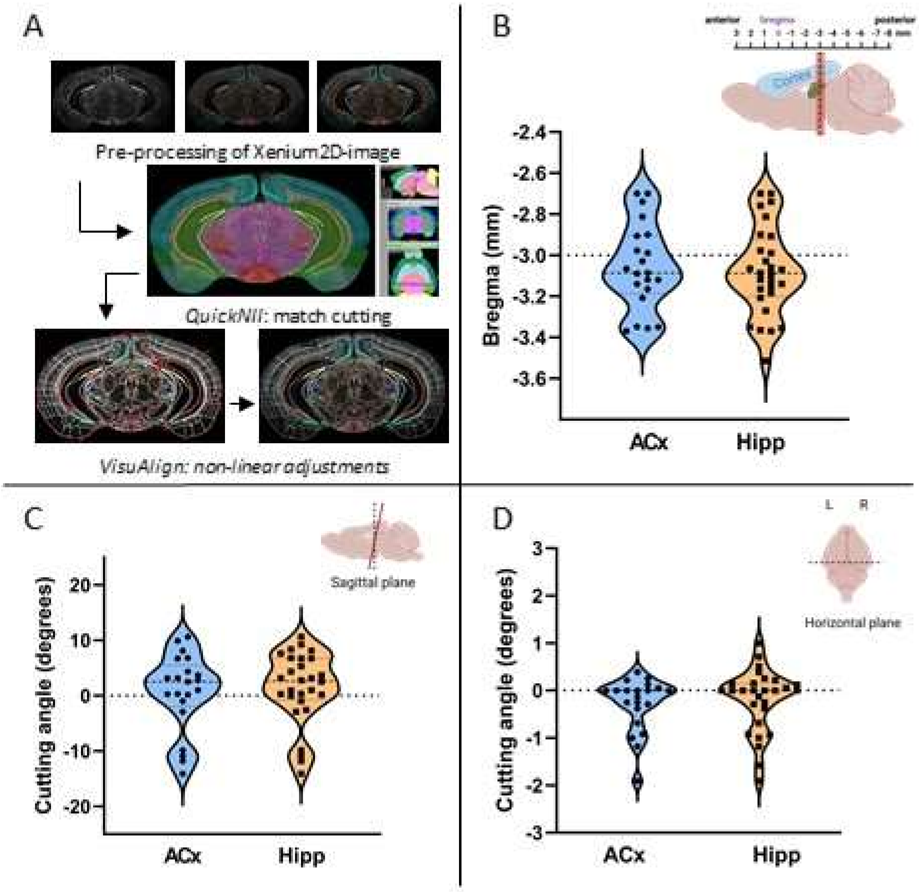


**Figure S1. Determining the positions and angles of coronal sections. A** Registration of sections to a reference atlas followed by non-linear adjustment. **B** Locations of the coronal sections on the anterior-posterior axis (target location -3.0 Bregma). **C** Cutting angles of the coronal sections in the sagittal plane. **D** Cutting angles of the coronal sections in the horizontal plane.

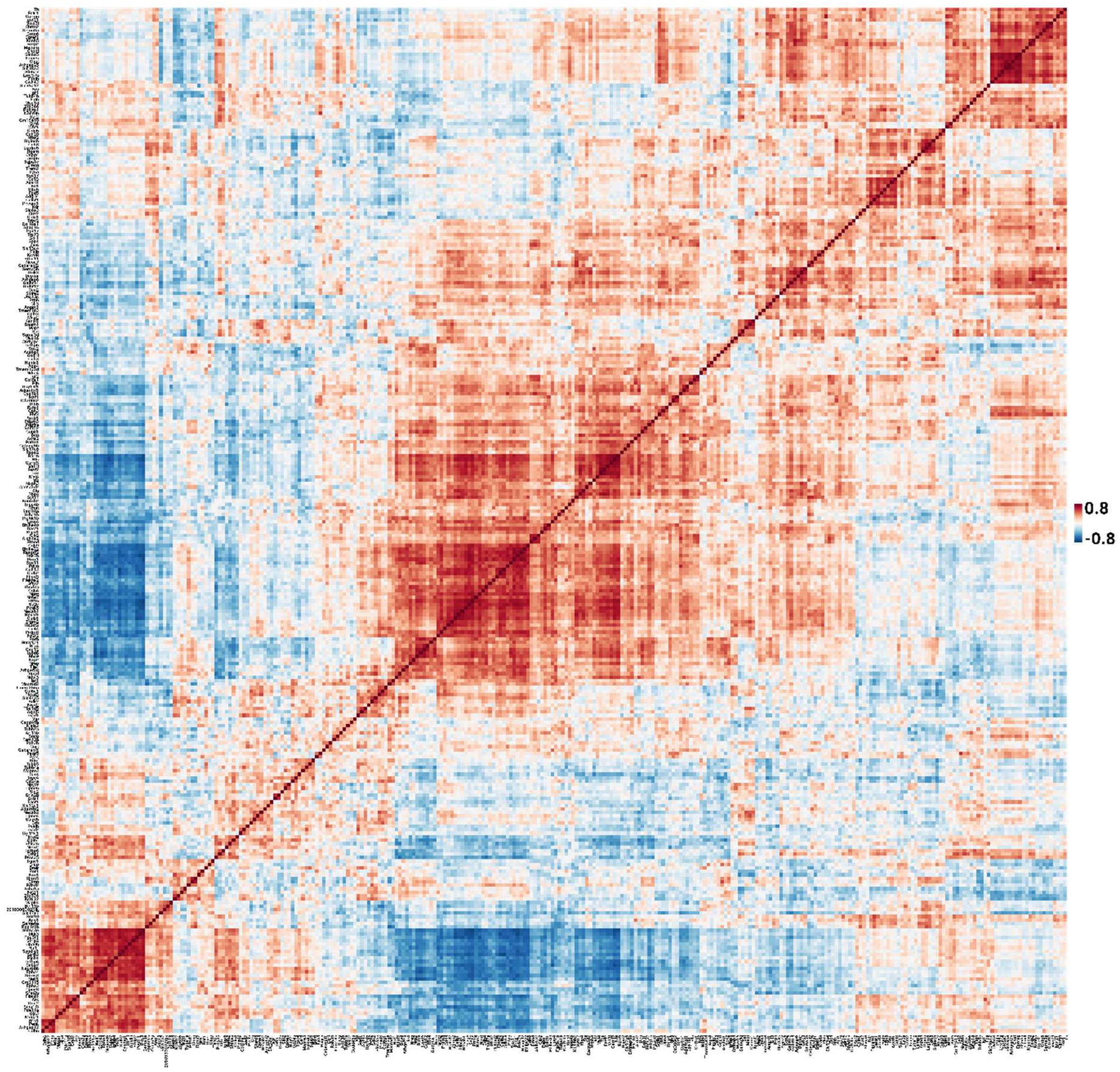


**Figure S2. Pairwise correlations (Pearson's r) among the 296 transcripts in the hippocampus.** Given the extensive correlation structure we ran principal component analysis for data reduction (see Figure S3).

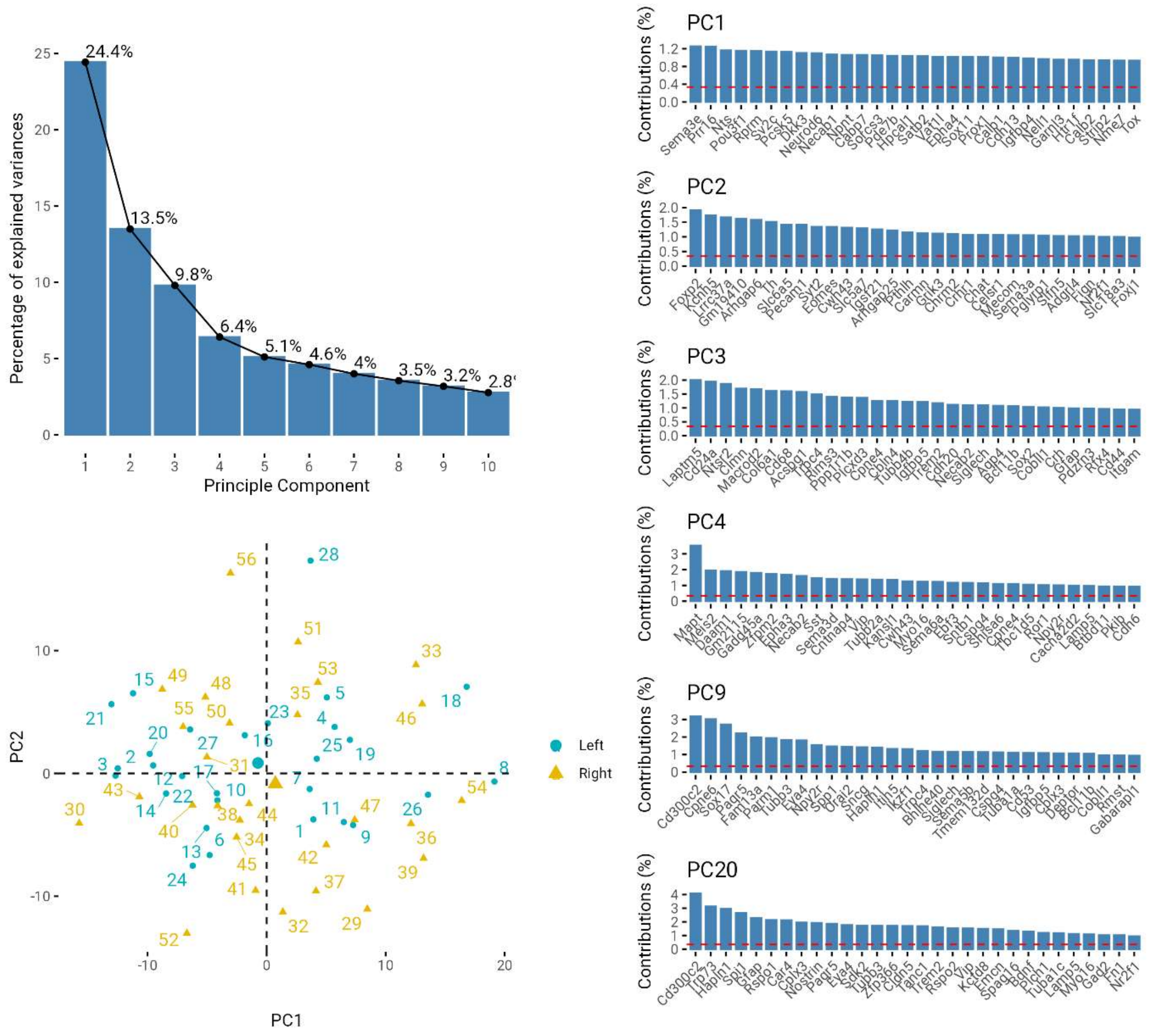


**Figure S3. Principal component (PC) analysis in the hippocampus.** The upper left panel illustrates the percentage of explained variance by the first ten PCs (18 PCs were needed to explain 90% of overall variance). The bottom left panel depicts the positions of sections and hemispheres in PC1-PC2 coordinates. The right panel shows the contributions (loadings) of the top 30 genes for six of the PCs, for illustration. Dashed red horizontal lines show the average loadings across 296 genes.

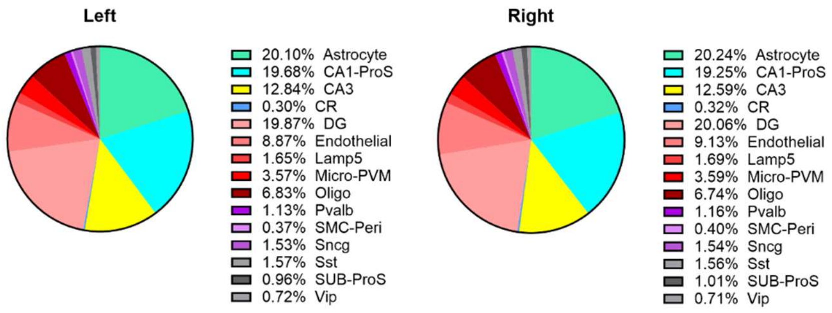


**Figure S4. Percentages of 15 major cell types in the left and right hippocampus.** All FDR adjusted P>0.35 when testing for hemispheric differences.

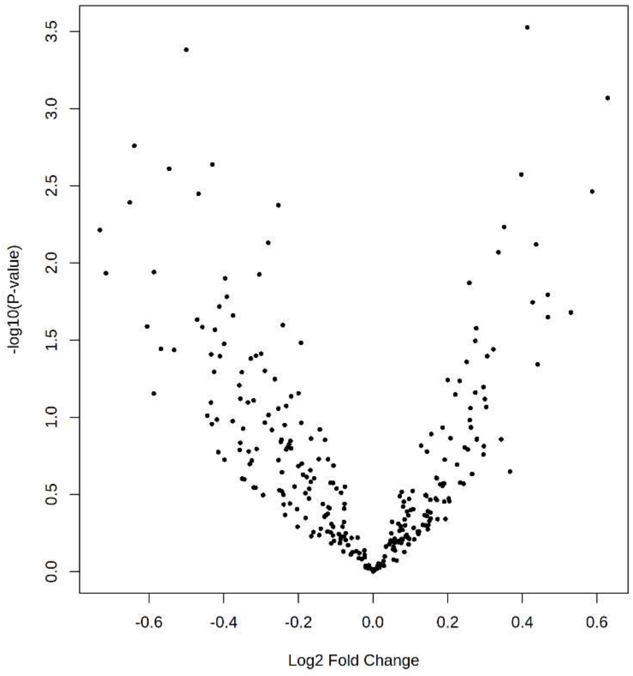


**Figure S5. Volcano plot of gene expression in the auditory cortex.** Each dot denotes a transcript. The x-axis represents the $log_2$ fold change in the right hemisphere compared to the left hemisphere. The y-axis depicts the $-log_{10}$ p values. All FDR adjusted P>0.061 (296 transcripts).

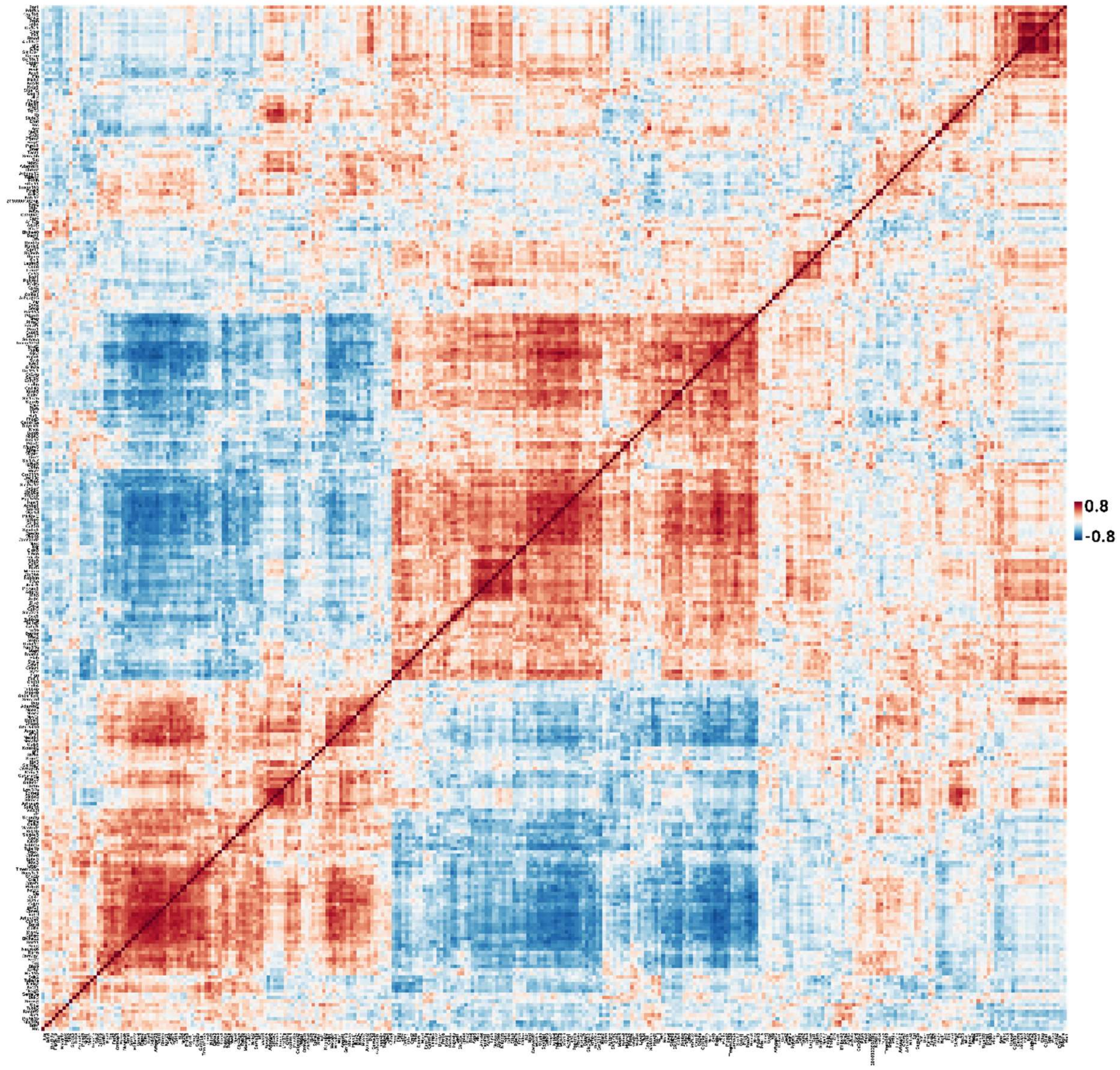


**Figure S6. Pairwise correlations (Pearson's r) among the 296 transcripts in the auditory cortex.** Given the extensive correlation structure we ran principal component analysis for data reduction (see Figure S7).

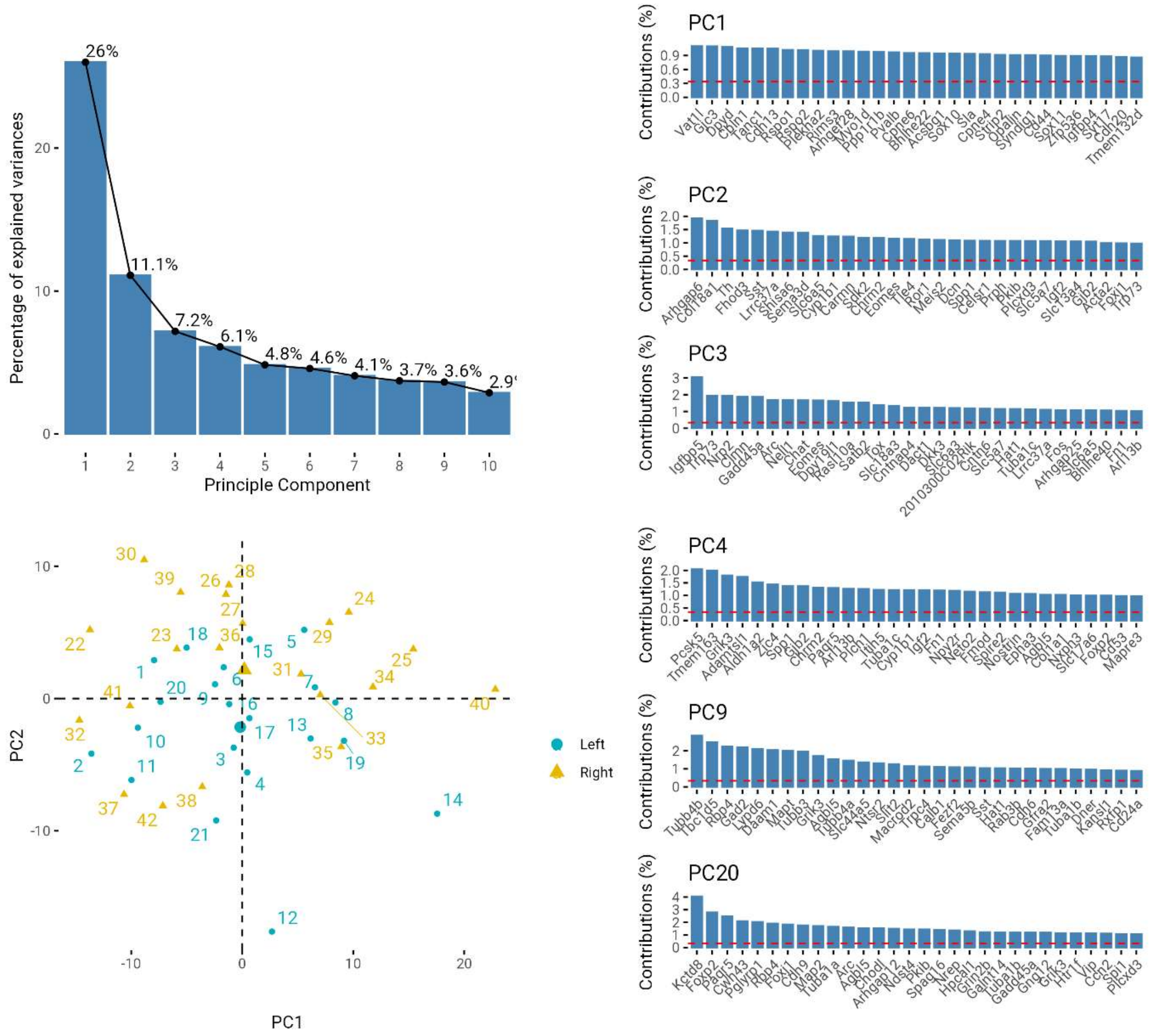


**Fig. S7 Principal component (PC) analysis in the auditory cortex.** The upper left panel illustrates the percentage of explained variance by the first ten PCs (20 PCs were needed to explain 90% of overall variance). The bottom left panel depicts the positions of sections and hemispheres in PC1-PC2 coordinates. The right panel shows the contributions (loadings) of the top 30 genes for six of the PCs, for illustration. Dashed red horizontal lines show the average loadings across 296 genes. PCs 2, 9 and 20 showed significant associations with hemisphere – see section 'Asymmetries in the auditory cortex' in the main text.
